# Handedness measures for the Human Connectome Project: Implications for data analysis

**DOI:** 10.1101/2020.03.08.982678

**Authors:** Lana Ruck, P. Thomas Schoenemann

## Abstract

Open data initiatives such as the UK Biobank and Human Connectome Project provide researchers with access to neuroimaging, genetic, and other data for large samples of left-and right-handed participants, allowing for more robust investigations of handedness than ever before. Handedness inventories are universal tools for assessing participant handedness in these large-scale neuroimaging contexts. These self-report measures are typically used to screen and recruit subjects, but they are also widely used as variables in statistical analyses of fMRI and other data. Recent investigations into the validity of handedness inventories, however, suggest that self-report data from these inventories might not reflect hand preference/performance as faithfully as previously thought. Using data from the Human Connectome Project, we assessed correspondence between three handedness measures – the Edinburgh Handedness Inventory (EHI), the Rolyan 9-hole pegboard, and grip strength – in 1179 healthy subjects. We show poor association between the different handedness measures, with roughly 10% of the sample having at least one behavioral measure which indicates hand-performance bias *opposite* to the EHI score, and over 65% of left-handers having one or more mismatched handedness scores. We discuss implications for future work, urging researchers to critically consider direction, degree, *and* consistency of handedness in their data.

## Introduction and motivation

Since it was first published in 1971, the Edinburgh Handedness Inventory (henceforth EHI; Oldfield, 1971) has become the gold-standard for evaluating handedness in neuropsychological contexts. The use of some version of the EHI is now ubiquitous for subject screening and participant exclusion in neuroimaging studies, often to the omission of other methods of handedness assessment (Fazio & Cantor, 2015). The exclusion of left-handed subjects (defined as those with either low or negative EHI scores) in neuroimaging contexts—specifically those probing language or other lateralized functions—is largely justified, considering the effect of subject lateralization on variance in activation patterns and resultant issues with common statistical approaches (see Bailey, McMillan, & Newman, 2019; Króliczak, Gonzalez, & Carey, 2019; Vingerhoets, 2014). Still, some have argued that this is a misguided approach, claiming that the inclusion of left-handers and other atypically lateralized individuals can tell us *more*, not less, about cortical function, and especially, asymmetry (Willems et al., 2014). Unfortunately, after decades of research paradigms which purposefully included left-handers, handedness effects on other aspects of laterality are still poorly understood. Efforts to identify a coherent relationship between handedness and language laterality, for example, have proceeded for over a century, and yet still, many studies show contrasting relationships between subject handedness and hemispheric activation patterns (Badzakova-Trajkov et al., 2016; Mazoyer et al., 2014; Mellet et al., 2014; Somers et al., 2015; Zago et al., 2016).

Disparate findings from large-scale studies of functional brain lateralization and handedness have resulted in revived discussions – at least from *within* the community of laterality researchers – on the use of the EHI and other handedness inventories. One result of these renewed discussions is a “degree vs. direction” approach, where subjects are classed not into binary directional categories (right- vs. left-hander), but into ordinal categories reflecting degree of handedness, such as strong right-handed, mixed-handed, weak left-handed, etc., prior to analysis (Gonzalez & Goodale, 2009; Kaploun & Abeare, 2010; Newman et al., 2014; Pritchard et al., 2013; Somers et al., 2015). This approach has led to some improvements, most notably in linking variability in left-handers’ hand performance, which is generally lacking in right-handers, to similar variability in activation patterns in the brain (Christman et al., 2015). Still, the partition of EHI scores into ordinal categories is arguably a least-effort and *post hoc* approach to addressing issues with handedness measures and classification, and – despite many critiques – the use of EHI values as a singular handedness assessment remains consistent.

Much was done to evaluate the internal consistency and validity of handedness inventories, and to compare various handedness assessments to each other, in the 1970’s and 1980’s. Overall, however, the expediency of self-report measures like the EHI resulted in their ubiquity over other handedness assessments (especially behavioral ones, which take time and resources to administer). Handedness literature from the past decade is arguably returning to these roots, as more recent discussions parallel earlier debates on handedness assessment and characterization (see Annett, 1985; Bryden, 1977; McManus, 1984, 1985, 1986). Despite a revived interest in the validity of handedness inventories (Büsch et al., 2010; Dragovic, 2004; Fazio et al., 2012, 2013; Milenkovich & Dragovic, 2013; Veale, 2014), to our knowledge, only a few studies have been conducted directly comparing self-report, survey-based handedness classifications with actual measures of hand performance and hand preference (Brown et al., 2004, 2006; Bryden et al., 2011; Corey et al., 2001; Flindall & Gonzalez, 2018; Gonzalez et al., 2007; Leppanen et al., 2018; McManus, Van Horn, & Bryden, 2016). Some of these studies evaluated a different handedness inventory as their survey-based measure (such as the Waterloo Handedness Questionnaire, or WHQ), but in general, participants in each of these studies were asked to complete both the survey *and* behavioral handedness assessments. These tasks typically include: at least one version of a pegboard task (Annett, grooved, etc.), finger tapping, grip strength, and other behavioral measures. Raw scores and laterality indices (or LI’s, see Methods below) for performance measures are then compared to the survey score, which is also a laterality quotient (see Edlin et al., 2015, however, for a discussion on EHI administration and scoring inconsistencies; also see McManus, Van Horn, & Bryden, 2016: 387-388, for a discussion on simple differences between hands versus laterality index calculations).

Brown and colleagues (2004) assessed the correlations between five performance measures (finger tapping, grip strength, both pegboard and grooved pegboard, and the Wathand Box Test) and the WHQ in 62 participants, and found that most measures had significant correspondence to the WHQ, with the exception of the grooved pegboard. They later (2006) conducted the same measures on 120 additional subjects, and found the same results – that all measures save for grip strength were significantly correlated with the WHQ values. However, in assessing directional correspondence between behavior-based scores and the WHQ (see Methods below), many subjects (ranging between 7% and 49%) showed at least one behavioral measure with a different handedness direction than their WHQ score (Brown et al., 2006: 8). A backward linear regression for predicting WHQ with behavioral scores showed the Wathand box test (WBT) as the most significant predictor of WHQ scores, and that grip strength was the only non-significant predictor. In a third study, they assessed the WBT and the WHQ as they related to language laterality in 142 subjects, using the Fused Dichotic Words Test (Bryden et al., 2011). They found that both tasks – one behavioral and one survey-based – had non-significant correlations with the language data, with the exception of the WHQ and language laterality in females. They conclude by urging further work comparing behavioral handedness measures to language lateralization, implicating fMRI as a component of future research.

Corey and colleagues (2001) assessed the correspondence between two performance measures including the EHI, and finger tapping, grip strength, and a grooved pegboard task, in 52 healthy subjects. They found high correspondence between preference and performance data, and only 5 subjects showed different handedness classifications between performance LI’s and survey classifications (two left-handers and 3 right-handers). Despite this high correspondence, they recommend the use of multiple metrics to classify subjects based on handedness. They conclude:

> “Whether other anatomic asymmetries are linked to hand preference or performance or both remains unclear. A multivariate approach to defining handedness and a multivariate examination of anatomic asymmetries may clarify the relationship of handedness to other more complex lateralized behaviors such as speech, language, and praxis”.
>
> — (Corey et al., 2001: 151)

Overall, these studies suggest that there are complicated patterns of association even between the different measures of handedness, and that survey-based and behavior-based handedness assessments are likely not isomorphic. These issues, obviously, would complicate any efforts to understand how handedness relates to other human asymmetries (see Gonzalez et al. 2018; Hopkins, 2018, for recent discussions of these topics).

In contrast to these approaches, Gonzalez and colleagues (2009) assessed correspondence between the EHI and hand-performance using more naturalistic tasks – puzzle and LEGO®-building – in 20 subjects. They filmed subjects’ hand movements and created LI’s for how often subjects used each hand, and then compared these data to EHI scores. They showed high correspondence between the LEGO® and puzzle tasks for all subjects, but these data did not match well with the EHI scores, especially for left-handers. Although all 10 left-handers in the sample were classed as strong left-handed (average EHI = –94.1), at least half of them used their right hand more frequently than their left in the behavioral task (Gonzalez et al., 2007: 277). In a later study probing language laterality with a dichotic listening task, they assessed correlations between the EHI, grip strength, finger tapping, and the LEGO® task in 36 subjects (Gonzalez & Goodale, 2009). They found that the LEGO® task measures were the only ones with significant correlations with the dichotic listening task. Although they also found significant correspondence between the EHI, grip strength scores, and finger tapping scores, these measures did not correlate with the dichotic listening task language laterality, as the LEGO® task did. They suggest that, in their sample, “there is something about visuomotor control and handedness that does not map onto other measures of laterality in motor control” such as the EHI and more common behavioral tasks (Gonzalez & Goodale, 2009: 3187).

In another recent paper (Flindall & Gonzalez, 2018), this team combined two handedness inventories, the EHI and Waterloo Handedness Questionnaire (for a combined survey, the E-WHQ) and assessed the survey’s accuracy, reliability, and ability to predict grasping patterns. In a meta-analysis of data from their previous studies, Flindall and Gonzalez (2018) showed that % right hand use for grasp-to-build tasks correlated significantly with E-WHQ survey scores, but only for the *entire* population. This effect was *not* replicated when subjects were grouped into subsets by handedness, as all correlations between E-WHQ scores and % right-hand use in Left, Right-, and Ambidextrous-handers were non-significant; they state that “within self-defined handedness groups, individual score on the E-WHQ is not useful in predicting right-hand preference in a simple grasping task” (Flindall & Gonzalez, 2018: 7). They also assessed whether or not the consistent Likert design of the survey impacted subject responses, showing that scrambling the response order for the 22-item survey led to significantly lower scores than the traditional test, within the same subjects. Finally, they assessed the consistency of subject responses in a test re-test paradigm, and found that over 90% of participants changed their response to at least one question. Overall, the team concludes that: “…the accuracy of a single handedness determination may be questionable; at worst, E-WHQ handedness scores may be irrelevant when it comes to predicting hand preference for grasping” (Flindall & Gonzalez, 2018: 13).

There seems to be little consensus on which measure is the best to use in explorations of handedness-related asymmetries, specifically for language, but extending into other functions as well. Although more naturalistic tasks may be the most viable option, they require more time and effort on the part of the experimenter in terms of recording, coding, and analyzing; this, we feel, explains the ubiquity of survey-based handedness assessments within neuropsychological research, and is one of the largest obstacles to overcome in future work. Aside from the use of more naturalistic manual motor tasks in the studies described above, relatively little has been done to explore the role that the EHI itself, and other handedness measures, may play in obscuring, not elucidating, the complexity of human handedness and its relationship with other phenomena. To add to this discourse, we assess multiple measures of handedness using handedness data from the Human Connectome Project (Van Essen et al., 2013).

## Methods

### About the HCP Data

The Human Connectome Project is an open-data initiative which provides structural and functional neuroimaging data to researchers (along with demographic data, and behavioral data for several common psychological assessments) from over 1200 healthy adult participants from the United States. All Human Connectome Project (HCP) subjects complete a battery of tasks, three of which are assessments of participant handedness^1^. The first is the 10-question, or “short form” version, of the Edinburgh Handedness Inventory (or EHI, see Oldfield, 1971), which is a standard self-reported survey of hand-use preference for various tasks, and the other two are hand motor measures from the NIH toolbox (Gershon et al., 2010; Kallen et al., 2012a, 2012b; Reuben et al., 2013; see also Wang et al., 2013). These measures include the 9-hole Pegboard Dexterity Test (or Rolyan pegboard) which measures time taken to complete a peg-manipulation task, in seconds (lower values reflect better performance); and a grip strength task, measured in pounds using a dynamometer (higher values mean better performance). Importantly, subjects complete the pegboard and grip strength tasks with both the left- and the right-hand, thus providing *behavioral* metrics for comparing the laterality of hand motor skills to the self-reported data from the EHI. In addition to these three metrics, the NIH toolbox includes a direct question: “Are you right-handed or left-handed?” – with possible responses being: “Right-handed” “Left-handed” and “Not sure” – so we treat these responses as a participant’s self-identified handedness. As subject recruitment was conducted irrespective of handedness, the number of left-handers in this sample, as classed by the EHI (*n* = 112, roughly 10%), corresponds well with broad handedness-trends across living human populations. Thus, we feel that this is a reliable sample for assessing correspondence between self-reported EHI values and direct physical measures of manual motor skill. Furthermore, with the impressive amount of behavioral and neuroimaging data also provided for these subjects (Barch et al., 2013), the HCP sample provides many future avenues for assessing handedness-related differences in psychometric data, brain anatomy, and brain function.

The HCP has released data for over 1200 subjects, but this sample includes several twin, sibling, and parent-child pairs. As we did not exclude for family or twin status in this study, it is important to note that roughly 300 of the 1200 HCP participants are related to at least one other participant. These data present a unique opportunity to study family relationships in future work, and indeed there are undoubtedly many interesting interactions to study between age, sex, twin status, and handedness in this data set. As effects of sex, age, and other demographics may be as related to the HCP’s recruitment and screening procedures as to a true signal, we will focus solely on the handedness measures in this paper, save for our discussion of the Freesurfer (brain) data. Of the 1200 HCP participants, those with positive illicit-drug test results (restricted-access^1^ demographic information) were first removed from the sample. Those with either the grip strength or pegboard measures three standard deviations above or below the sample mean were also removed from analysis. Thus, this study includes 1179 of the 1200 HCP subjects. Some of these 1179 participants did not undergo neuroimaging, and do not have Freesurfer data (see the section on *Subject classification in neuroimaging contexts*), so our sample for the Freesurfer data is 1096 participants. In addition, 46 HCP participants underwent a full test-retest paradigm and have two sets of data to compare for internal consistency of the HCP protocols; one of the test-retest participants was stripped from the larger sample due to outlying handedness scores, so our test-rest data includes 45 participants.

### Measures and terminology

Our main goal is to explore the typical ways in which neuroimagers partition left- and right-handers into groups for analysis. Thus, many of our analyses are run on the entire HCP sample as well as on the right- and left-handed subject subsets, *as defined by EHI scores*, to mirror the approach of others working with handedness data in neuroimaging contexts. Analyses done on EHI-delineated ‘right-handers’ (EHI > 0) will be indicated by the term **EHI+** and those on EHI-delineated ‘left-handers’ (EHI < 0) will be denoted **EHI–**. To reiterate, these terms reflect subject groupings based on EHI scores, and are independent of the behavioral handedness measures for the right- or left-hand, and of the self-identification groupings, which will be indicated as such. We report summary statistics on the EHI and raw grip strength and pegboard scores across all 1179 subjects, as well as ‘left-handers’ (EHI– *n* = 112) and ‘right-handers’ (EHI+ *n* = 1067) separately, in Table 1. Based on the self-identification data alone, discrepancies exist between handedness classification based on EHI scores and subject self-reporting: 120 subjects self-identified as left-handed, 1045 self-identified as right-handed, and 14 self-identified as not sure. As with previous studies (Flindall & Gonzalez, 2018; Mazoyer et al., 2014), several (*n* = 25) HCP participants with “moderately” right-handed EHI scores (EHI+ participants) self-identified as *left handed*, and a small number (*n =* 3) of participants with left-handed EHI scores (EHI– participants) identified as *right-handed* in the HCP sample as well.

**Table 1:** Summary statistics for the entire subject pool (top, *n* = 1179), and the right-hander (middle) and left-hander (bottom) subsets. Mean, standard error (s.e.), population variance, standard deviation (st.dev.) are shown for all groups for each of the following measures: raw Grip Strength (grip strength) scores for the right- and left-hands; raw Pegboard (pegboard) times (in seconds) for the right- and left-hands; EHI survey scores; and calculated Grip Strength and Pegboard Laterality Indices (grip strength LI, pegboard LI).

| Summary Statistics |  | Grip strength right hand (pounds) | Grip strength left hand (pounds) | Pegboard right hand (seconds) | Pegboard left hand (seconds) | EHI | Grip strength LI | Pegboard LI |
| --- | --- | --- | --- | --- | --- | --- | --- | --- |
| All subjects<br>n = 1179 | mean | 86.531 | 81.167 | 21.630 | 22.604 | 66.068 | 3.426 | 2.270 |
|  | s.e. | 0.781 | 0.770 | 0.083 | 0.078 | 1.301 | 0.173 | 0.173 |
|  | variance | 26.804 | 26.443 | 2.839 | 2.667 | 1993.01 | 35.444 | 35.549 |
|  | st.dev. | 717.833 | 698.657 | 8.055 | 7.108 | 44.662 | 5.956 | 5.964 |
| EHI+<br>n = 1067 | mean | 86.242 | 80.403 | 21.521 | 22.686 | 78.932 | 3.780 | 2.699 |
|  | s.e. | 0.817 | 0.810 | 0.086 | 0.082 | 0.593 | 0.179 | 0.179 |
|  | variance | 26.686 | 26.472 | 2.821 | 2.681 | 374.913 | 34.295 | 34.321 |
|  | st.dev. | 711.475 | 700.086 | 7.953 | 7.179 | 19.372 | 5.859 | 5.861 |
| EHI-<br>n = 112 | mean | 89.282 | 88.439 | 22.670 | 21.827 | -56.473 | 0.061 | -1.814 |
|  | s.e. | 2.634 | 2.376 | 0.266 | 0.228 | 2.710 | 0.552 | 0.510 |
|  | variance | 27.874 | 25.144 | 2.812 | 2.411 | 815.464 | 33.881 | 28.828 |
|  | st.dev. | 770.037 | 626.591 | 7.836 | 5.761 | 28.685 | 5.847 | 5.393 |

Due to our particular interest in whether the EHI accurately reflects handedness when measured in other domains, we calculated laterality indices (henceforth, LI or LI’s) for the behavioral tasks, using the following approach:

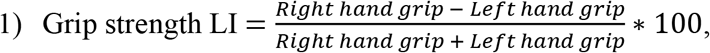

where positive values indicate a right-hand superiority, and negative values indicate a leftward bias.

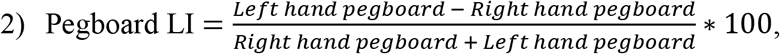

where positive values indicate a right-hand superiority, and negative values indicate a leftward bias (recall that pegboard scores are in seconds, so lower measures reflect better performance).

These LI’s preserve directional bias within subjects in a way similar to the EHI—positive values indicate rightward bias; negative values indicate leftward bias—and they also account for absolute differences (i.e., magnitude differences in raw scores) between subjects (Brown et al., 2006; Oldfield, 1971; but see McManus, Van Horn, and Bryden, 2016: 387-389). Discrepancies between the EHI and these other handedness measures are the focus of the remaining sections.

## Results

### Group-level differences in HCP handedness measures

As with other data sets, the HCP behavioral handedness measures do not have the classic “j-shaped” skew which is present in EHI scores; grip strength and pegboard LI values are generally centered around 0 and are more normally distributed, particularly in the case of EHI– subjects. Although many of the HCP handedness measures are non-normally distributed, we use parametric statistics in this study following the recommendation of Fagerland (2012), in which parametric statistics were shown to be more appropriate for large-scale data sets, even in cases where the distributions are skewed. All analyses were completed in R 3.5.1 (R core Team 2018). We used t-tests to assess group-level differences in the raw NIH Toolbox measures, and all measures except for right-hand grip strength showed significant differences between EHI+ and EHI– subjects (see Table 2).

**Table 2:** *t*-tests comparing right-handed subset (EHI+) means to left-handed subset (EHI–) means, for: raw grip strength for the right- and left-hands; raw pegboard times (in seconds) for the right- and left-hands; EHI values; and calculated grip strength and pegboard LI’s. Means, difference in means, t-values, p-values, and effect sizes are shown. * *p* < 0.05 ** *p* < 0.01 *** *p* < 0.001

| ***t*-tests for EHI-based handedness groups** | **EHI+ mean** | **EHI– mean** | **Difference in means** | ***t*-value** | ***p*-value** | **Cohen's D (effect size)** |
| --- | --- | --- | --- | --- | --- | --- |
| **grip strength right hand** | 86.242 | 89.282 | -3.040 | 1.102 | 0.2723 | 0.113 |
| **grip strength left hand** | 80.403 | 88.439 | -8.036 | 3.201 | 0.002** | 0.305 |
| **pegboard right hand** | 21.521 | 22.670 | -1.149 | 4.175 | <0.001*** | 0.419 |
| **pegboard left hand** | 22.686 | 21.827 | 0.859 | -3.429 | <0.001*** | 0.323 |
| **EHI** | 78.932 | -56.473 | 135.405 | -48.80 | <0.001*** | 6.627 |
| **Grip Strength LI** | 3.780 | 0.061 | 3.719 | -6.403 | <0.001*** | 0.635 |
| **Pegboard LI** | 2.699 | -1.814 | 4.513 | -8.327 | <0.001*** | 0.770 |

It is often the case that significant differences in handedness measures within un-balanced samples are driven largely by the ‘right-handed’ participants (see Flindall & Gonzalez, 2018: 8, Figure 1). Thus, performance differences between the dominant and non-dominant hands were tested within each handedness group. EHI+ subjects showed significantly better performance for the dominant hand compared to the non-dominant hand for both grip strength and pegboard (grip strength difference in means = 9.75, *t* = 6.451, *p* < 0.001, effect size = 0.385; pegboard difference in means = −2.54, *t* = −16.549, *p* < 0.001, effect size = 0.993), but EHI– subjects showed a significant difference *only* for the pegboard task (grip strength difference in means = −6.203, *t =* −1.249, *p* = 0.216, effect size = 0.291; pegboard difference in means = 1.963, *t* = 3.740, *p* < 0.001, effect size = 0.869) (see Figure 1). As shown in Table 2, effect sizes between EHI+ and EHI– participants vary widely between the EHI and the behavioral measures. Pearson’s linear correlations between the EHI and behavioral LI’s show small, but significant positive correlations between the three measures, indicating that increased rightward-bias in one measure correlates with increased rightward-bias in the others. Sub-sample tests, however, confirmed that this effect was *also* driven by the EHI+ subjects, as only one of the EHI+ correlations reached significance, and *none* of the EHI– correlations did (Table 3). Although we replicate group-level patterns typically reported in large-scale studies on handedness measures, our correlations suggest that EHI scores may be poor representations of HCP subjects’ actual manual performance, particularly for presumed ‘left-handers.’ These results corroborate a common characterization of left-handers in the broader literature, namely, that they are not left-biased in the same way that right handers are right-biased. This provides further justification for exploring left- and right-handed subjects as separate subsets (EHI+ and EHI–) in addition to running whole-group analyses.

**Figure 1:**
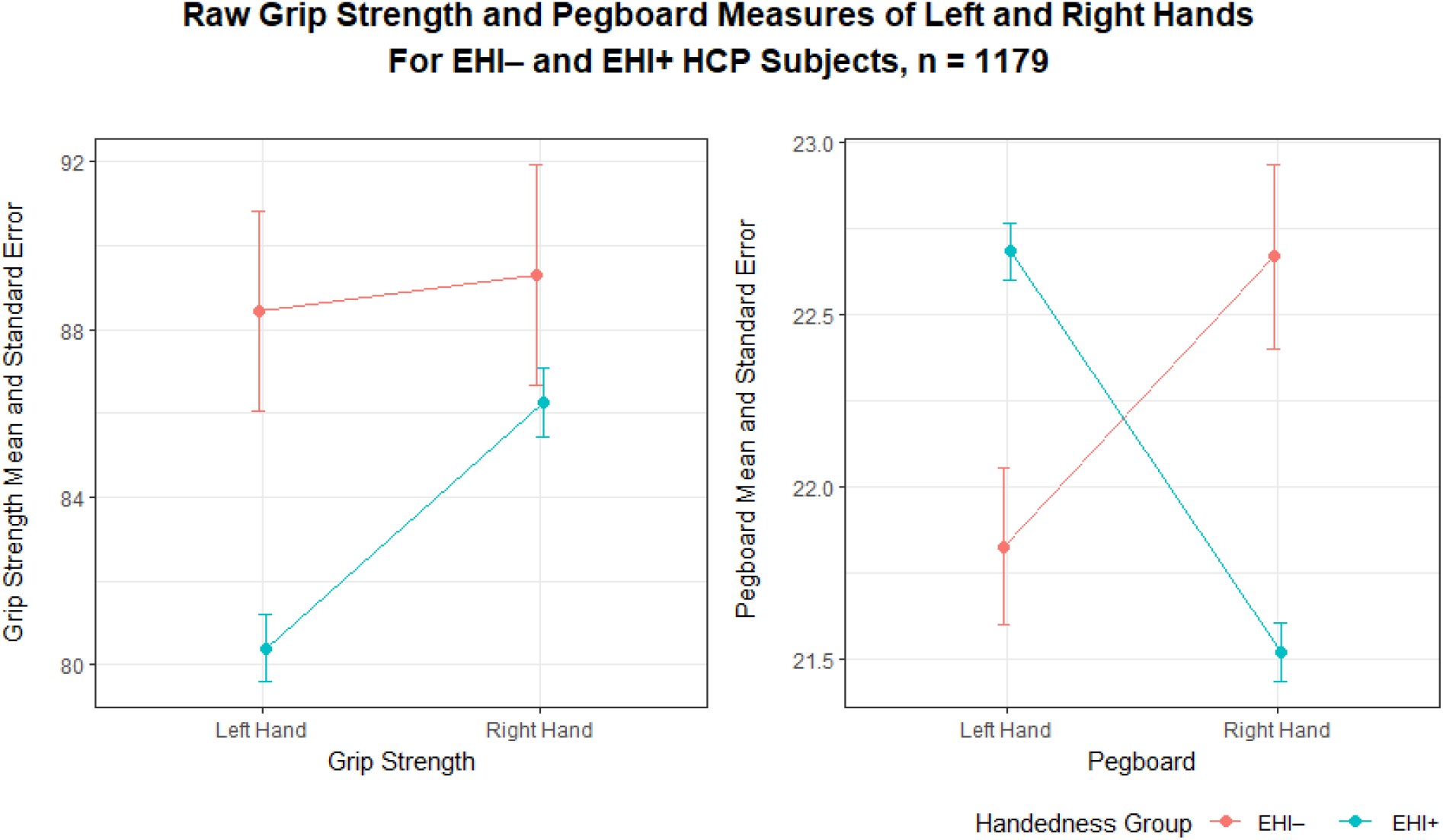
Interaction plots depicting raw grip strength (left) and raw pegboard (right) scores for EHI– (red) and EHI+ (aqua) subject subsets. Note that differences between the dominant and non-dominant hand are less severe in EHI– participants than in EHI+ ones, especially for grip strength.

**Table 3:** Pearson’s linear correlation values for EHI and grip strength LI, EHI and pegboard LI, and grip strength and pegboard LI’s. Correlation values (r) and probabilities (p(uncorrelated)) are shown for all three tests for the entire sample (top), as well as the EHI+ subset (middle) and EHI– subset (bottom). * *p* < 0.05 ** *p* < 0.01 *** *p* < 0.001

| Pearson's correlations for EHI groups | EHI and grip strength LI correlation (r) | p-value | EHI and pegboard LI correlation (r) | p-value | Grip strength and pegboard LI correlation (r) | p-value |
| --- | --- | --- | --- | --- | --- | --- |
| All subjects | 0.1866 | <0.001*** | 0.2511 | <0.001*** | 0.0848 | 0.003** |
| EHI+ | 0.0443 | 0.15 | 0.1452 | <0.001*** | 0.0515 | 0.09 |
| EHI- | 0.1119 | 0.24 | -0.0406 | 0.67 | -0.0107 | 0.91 |

### Characterizing handedness distributions: What about consistency?

That left- and right-handers show different patterns hand biases across multiple measures is interesting on its own, but that these differences are unstable across multiple measures complicates interpretations. An important discussion related to this about handedness *consistency* (see Prichard, Propper, & Christman, 2016), which is related to degree of preference. On this, Leppanen and colleagues (2018) state:

> “If handedness inventories are valid measures of preference, then reports of strong preference should be associated with relatively large performance disparities favoring the putatively preferred hand and reports of weak preference should be associated with relatively small disparities. To our knowledge, such a relationship has *never* been documented in the context of performing laboratory analogues of inventory tasks”.
>
> — (Leppanen et al., 2018: 544, emphasis added)

This team investigated 129 participants (only 8 with left-biased EHI scores), and hypothesized that EHI self-report responses would correlate with hand choice when participants actually performed the survey tasks spontaneously (untimed), and with time differences between the dominant and non-dominant hand when participants performed the same survey tasks again with both hands (timed). They found a significant correlation between EHI scores and hand choice for spontaneous task completion. Using a simple difference (non-dominant hand time – dominant hand time) for the timed version of each task, the team first noted that differences between hands were significantly larger when the non-preferred hand was used first (this was randomly assigned for each participant). As hand order was significant, the team did analyses for preferred-hand-first and preferred-hand-second groups separately; they found significant correlations between EHI scores and the magnitude of time difference between hands when performing survey tasks. Regarding handedness consistency, Leppanen and colleagues split their data set into consistent (EHI +/– 80) and inconsistent (EHI between 0 and +/–75) handers, and found that proportion of tasks performed with the preferred hand, and time differences between hands, were significantly higher in consistent handers, with the odds of using the nonpreferred hand for spontaneous task performance 8 times higher in inconsistent handers (Leppanen et al., 2018: 551).

We wanted to explore the HCP data in a similar way, so we split EHI+ and EHI– participants by handedness consistency using the same EHI cutoff (scores +/– 80 are considered consistent handers, whereas inconsistent handers have EHI scores between 0 and +/– 75). It is important to note that a significantly higher proportion of EHI+ participants are consistent handers (*n* = 662, 62.04%) when compared to EHI– participants (*n* = 30, 26.78% classed as consistent; **χ**-squared for EHI and consistency = 50.527, *p* < 0.001), again likely reflecting left-hander’s overall tendency towards reduced manual bias. The only measure which showed significant differences between consistent and inconsistent handers was the pegboard LI, and this was only in EHI+ participants (difference of means = 1.3803, *t* = 3.7157, *p* < 0.001, effect size = 0.238). Unlike Leppanen and colleagues (2018), and at a much larger sample size, we find that consistently-handed HCP participants have similar distributions of all other behavioral handedness measures, regardless of whether they are left-handed or right-handed (Figure 2).

**Figure 2:**
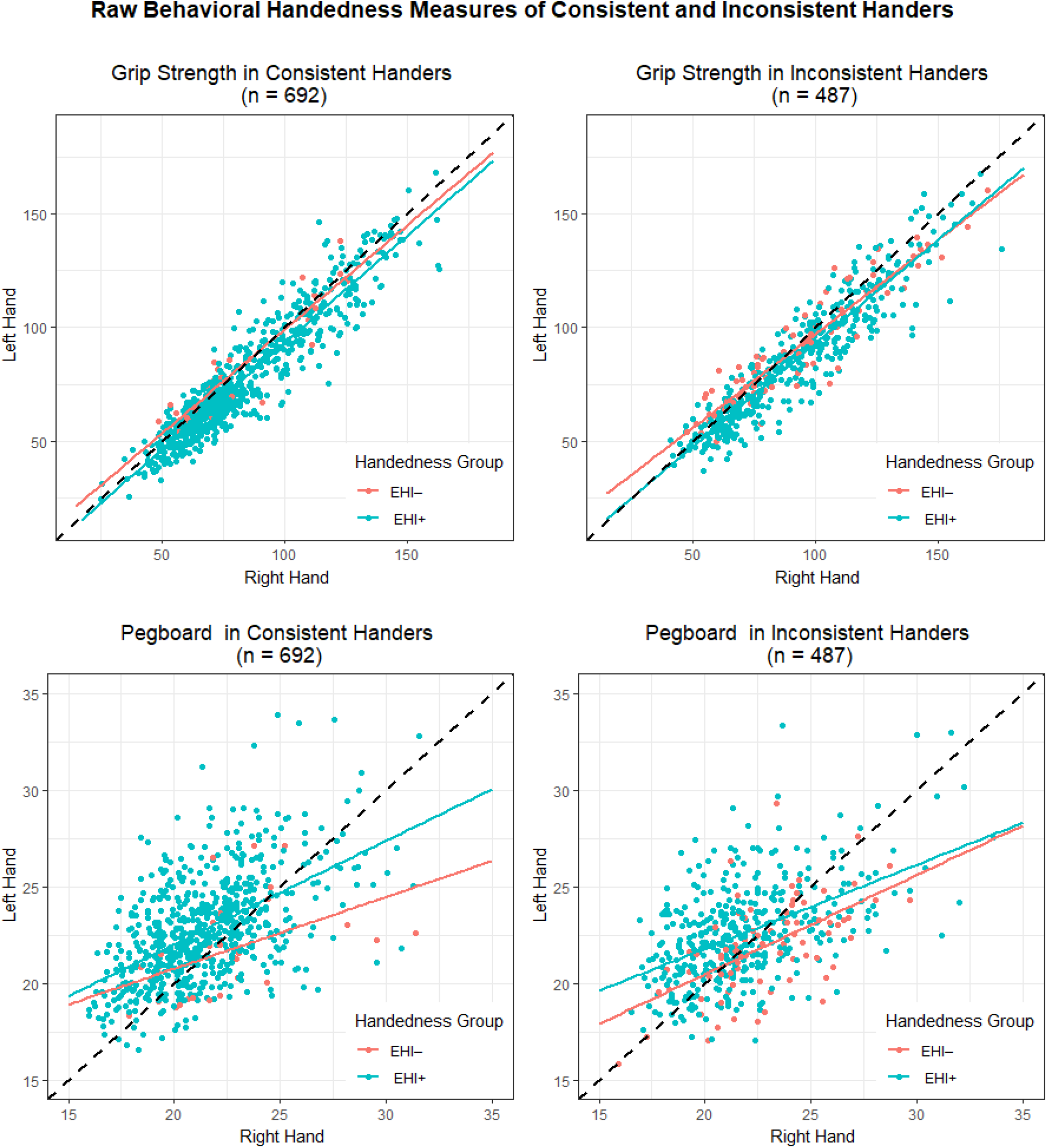
Raw scores for grip strength and pegboard in consistent vs. inconsistent handers. With the exception of pegboard scores, consistent and inconsistent handers (based on EHI scores) have indistinguishable behavioral handedness measures.

### Correspondence between the EHI and behavioral handedness measures

We have outlined several ways in which EHI scores *seem* useful for partitioning HCP subjects into meaningful handedness groups, but when evaluated against the behavioral measures, these procedures look more tenuous, which is concerning considering their ubiquity. McManus, Van Horn, and Bryden (2016) discussing related issues with regard to handedness characterization, revisiting data from Tapley and Bryden’s (1985) circle-marking task (*n* = 1556, with *n* = 161 self-identified as left-handed), and Van Horn’s (1992) study, which includes Tapley-Bryden circle-marking data as well as Annett pegboard data (where various pegboard configurations were tested). Although this team was not specifically interested in validating the EHI survey against behavioral measures, but instead in the overall characterization of handedness, they note “almost perfect separation of self-reported right and left-handers” on the Tapley-Bryden circle-marking task, with “only 7 (0.5%) right-handers performing better with the left hand, and 1 (0.6%) left-handers performing better with the right hand” (McManus, Van Horn, & Bryden, 2016: 378). McManus and colleagues (2016) found that direction of handedness across tasks was more consistent than degree of handedness, and note:

> “*If a person is right-handed for task A then they are very likely to be right-handed for task B, but if their dominant hand is very much better than their non-dominant hand for task A then that has no predictive ability for how much better their dominant hand will be com-pared with their non-dominant hand on task B.*”.
>
> — (McManus, Van Horn, & Bryden, 2016: 393, emphasis original)

We also assessed *congruency* between the three handedness scores for each subject. Does the direction of the EHI (positive for ‘right-handers’ and negative for ‘left-handers’) match the direction of the behavioral LI’s, which, as calculated, also have positive values to indicate rightward bias and negative values to indicate leftward bias? We considered congruency on a subject-specific basis using the following categories: full congruency, where all three scores indicate the same directional bias (all positive, or all negative); partial congruency, where one of the behavioral LI’s is in the opposite direction as the EHI; and non-congruency, where *both* the grip strength LI and pegboard LI indicate hand bias opposite to the EHI score. Congruency was assessed for the whole sample as well as the EHI+ and EHI– subsets (Table 4).

**Table 4:** Congruency of EHI direction with both Grip Strength (grip strength LI) and Pegboard (pegboard LI) for all subjects (top), the EHI+ subset (middle), and the EHI– subset (bottom). Congruency was assessed for each subject, where positive values indicate right-hand skew and negative values indicate left-hand skew for all three measures. Frequencies for congruent subjects (all measures match), partial congruence (EHI matches either grip strength LI or pegboard LI), and non-congruent (EHI does not match either grip strength LI or pegboard LI), are shown in the left, and percentages on the right. Note that roughly half of the overall sample has at least one behavioral score indicating opposing hand preference to the EHI, and more than 15% of left-handers (EHI–) have behavioral scores which both indicate right-hand preference.

| **Congruence** | **Total** | **Congruent** | **Partial** | **Incongruent** | **% Congruent** | **% Partial** | **% Incongruent** |
| --- | --- | --- | --- | --- | --- | --- | --- |
| **All subjects** | 1179 | 592 | 469 | 118 | 50.25 | 39.81 | 10.02 |
| **EHI+** | 1067 | 555 | 413 | 99 | 52.01 | 38.71 | 9.28 |
| **EHI–** | 112 | 37 | 56 | 19 | 33.04 | 50.00 | 16.96 |

Only half of all HCP subjects show full EHI–grip strength LI–pegboard LI congruency, although again, as with the raw scores, this trend is largely being driven by the EHI+ subjects in the sample (Figure 3). *Over 65%* of the left-handed subjects, as classed by the EHI, have at least one behavioral score which indicates a right-hand bias. Alarmingly, over 10% of HCP participants have behavioral LI’s which *both* indicate hand performance opposite to their EHI scores, regardless of whether they report as right-handed or left-handed. Intra-class correlations (ICC’s) for EHI, grip strength LI, and pegboard LI directionality (Koo et al., 2016) show low correspondence (< 0.2, indicating only slight agreement) for the whole sample, and for the EHI+ and EHI – subsets as well, confirming our other analyses (see Supplementary Materials, Table 1). Interestingly, congruence frequencies are similar when the data are further split by handedness consistency (see previous section), so even those with extreme EHI values have high incidences of incongruent behavioral data.

**Figure 3:**
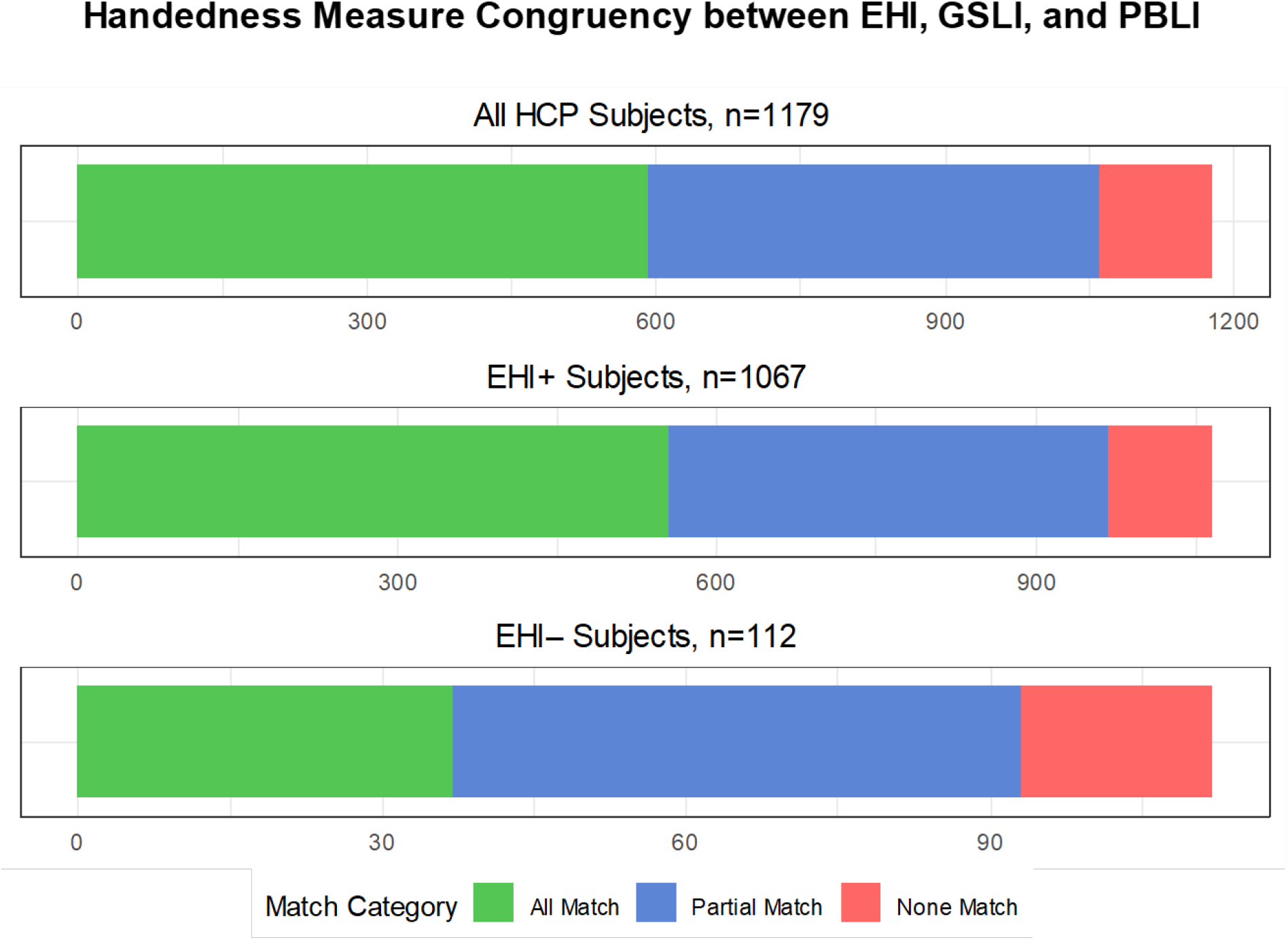
Congruency plots for the entire sample (left), EHI– (center) and EHI+ (right) subjects. A minority of EHI– subjects have all three scores matching in direction, and roughly half of all subjects have either grip strength LI or pegboard LI scores indicating hand performance bias opposite in direction to their EHI scores.

Discussing the Van Horn (1988) data, McManus and colleagues (2016) note that pegboard data are unimodal, as opposed to both the circle marking task (which is bimodal) and survey data (which is “j-shaped”). On this, they claim that “the T&B [circle] task separates the right- and left-handers entirely, whereas the pegboard scores *are not so good at doing that*, four of the 28 right-handers and 2 of the 28 left-handers being in the “wrong” half” (McManus, Van Horn, & Bryden, 2016: 384, emphasis added). We have found very different proportions of congruency in the HCP data, although, if anything, this confirms the authors original point that different handedness measures simply behave differently.

### Measurement reliability: HCP test-retest data

Unfortunately, knowing all the ways in which multiple handedness measures do not correspond with each other does very little to tell researchers which measure might be the best one to use in analyses about handedness. Of potential relevance here is the HCP’s test-retest data^2^, in which EHI, grip strength, and pegboard measures were collected on two separate occasions for 45 participants. Test-retest reliability was computed using Pearson’s correlations between initial test scores and retest scores (Table 5). Regarding self-report and survey-based EHI data, we replicate Flindall and Gonzalez’s (2018) findings that survey scores are generally stable across testing sessions, with the exception of the Broom question (“Which hand would you use to hold a broom (upper hand)?”), and Eye question (“Which eye do you use when using only one?”). Raw grip strength measures have high reliability as well, although raw pegboard scores have only moderate reliability. Presumably, measures with higher test-retest reliability should be favored, as low reliability likely indicates high measurement error. Despite high and moderate reliability at the raw score level, reliability scores for grip strength and pegboard *laterality indices* (or for raw differences; see McManus, Van Horn, & Bryden, 2016: 387-388) are much lower. It is unclear to us why this might be, but overall, it seems that as one looks at handedness data for these large-scale data sets more thoroughly, there is less and less clarity about how they should best be used.

**Table 5:**
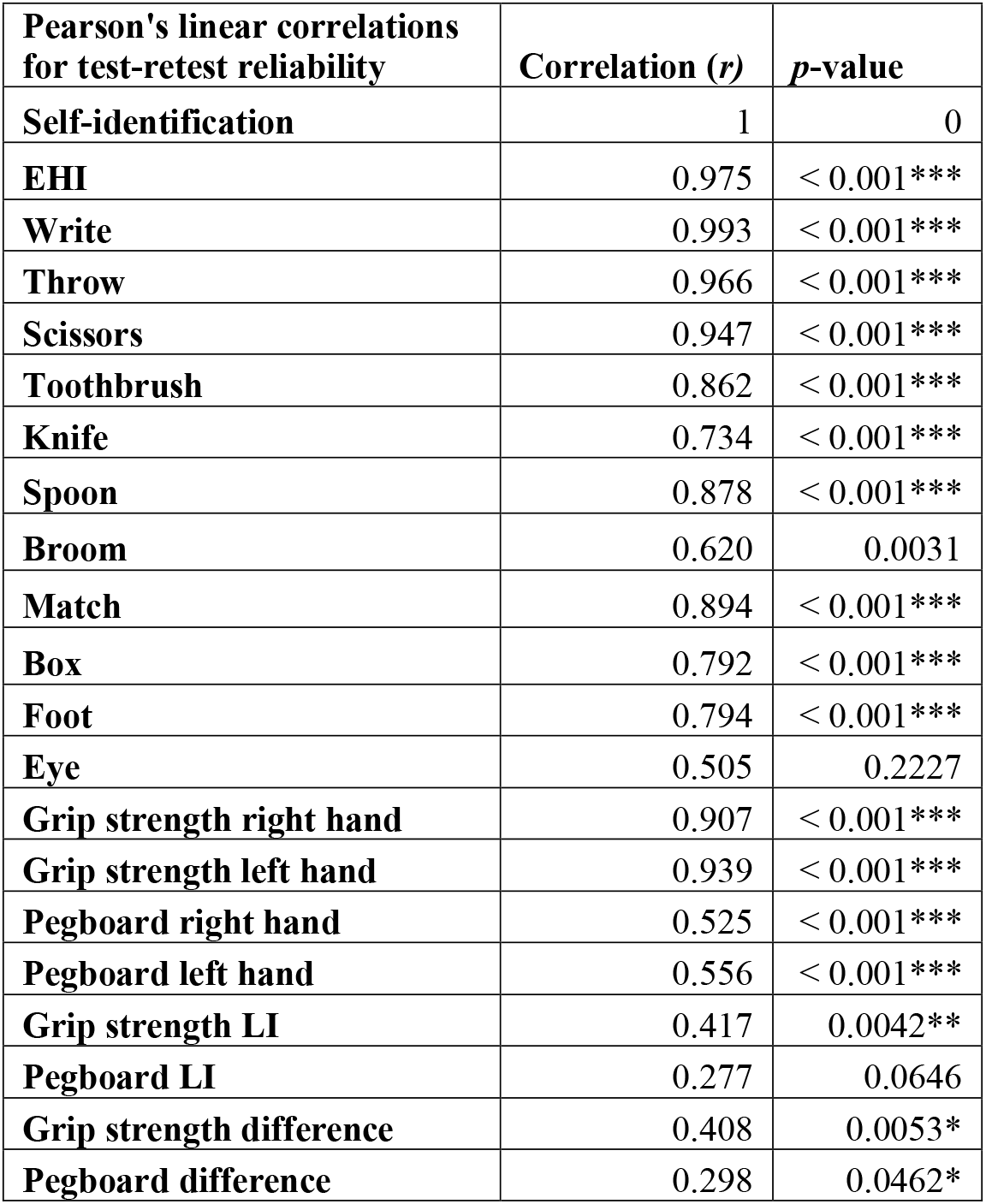
Test-retest reliability measures, using Pearson’s linear correlations, for the entire test-retest sample (*n* = 45). Correlation values (*r*) and probabilities (*p*(uncorrelated)) are shown for self-identification, overall EHI scores, the individual survey question answers, raw grip strength and pegboard scores, and laterality indices and raw differences between these measures for the right and left hands. * *p* < 0.05 ** *p* < 0.01 *** *p* < 0.001

#### A multivariate approach for aggregating handedness scores

Although our analyses of these three handedness metrics are interesting in their own right, we are ultimately less interested in predicting handedness from one measure to the next, and are instead more concerned with implications for subject classification and exclusion in neuroimaging contexts. Knowing that roughly half of the HCP subjects have partially or entirely mismatched survey-based and behavior-based handedness scores, and that each of these measures is genuinely complex in terms of its reliability and potential external validity, what procedures should be used to split subjects into groups for analysis? What, if anything, should be done about the use of the EHI for subject recruitment and screening? Results of many studies could change if subjects were grouped as ‘right-’ or ‘left-handed’ based not on EHI scores, but on Grip Strength (grip strength LI+ *n* = 845 (71.7%); grip strength LI– *n* = 334 (29.3%)), Pegboard (pegboard LI+ *n* = 774 (65.6%); pegboard LI– *n* = 405 (35.4%)), or other handedness measures. Unfortunately, so little has been done to assess these measures in their own right that much more needs to be done to assess their validity if we are to justify their use *in place* of the EHI. In lieu of this, we advocate for a multivariate approach (following Corey et al., 2001) by applying a Principal Component Analysis to the EHI, grip strength LI and pegboard LI scores together. Principal Component Analysis (PCA) is a widely used statistical approach for assessing the degree to which multiple variables can (or cannot) be reduced to a smaller number of dimensions (Wold, Esbensen, & Geladi, 1987). PCA analysis including the EHI, grip strength LI, and pegboard LI, resulted in three components (PC’s, see Table 6): PC1 (45.19% variance explained) has positive loadings on all three handedness measures, and we suggest it as a good measure of **overall handedness** (with negative scores reflecting overall left-hand bias, positive values reflecting overall right-hand bias), and intermediate values reflecting inconsistent or mixed handedness (Figure 4). PC2 (30.67% variance explained) appears to represent a dexterity vs. strength tradeoff (positive grip strength LI loading, negative pegboard LI loading), and PC3 (24.14% variance explained) contrasts the EHI (self-reported) against the behavioral handedness measures.

**Table 6:** Loadings for EHI, grip strength LI, and pegboard LI on PC1, PC2, and PC3.

| PCA Loadings | PC1 | PC2 | PC3 |
| --- | --- | --- | --- |
| EHI | 0.659 | -0.088 | 0.747 |
| Grip strength LI | 0.577 | 0.812 | -0.330 |
| Pegboard LI | 0.482 | -0.578 | -0.577 |

**Figure 4:**
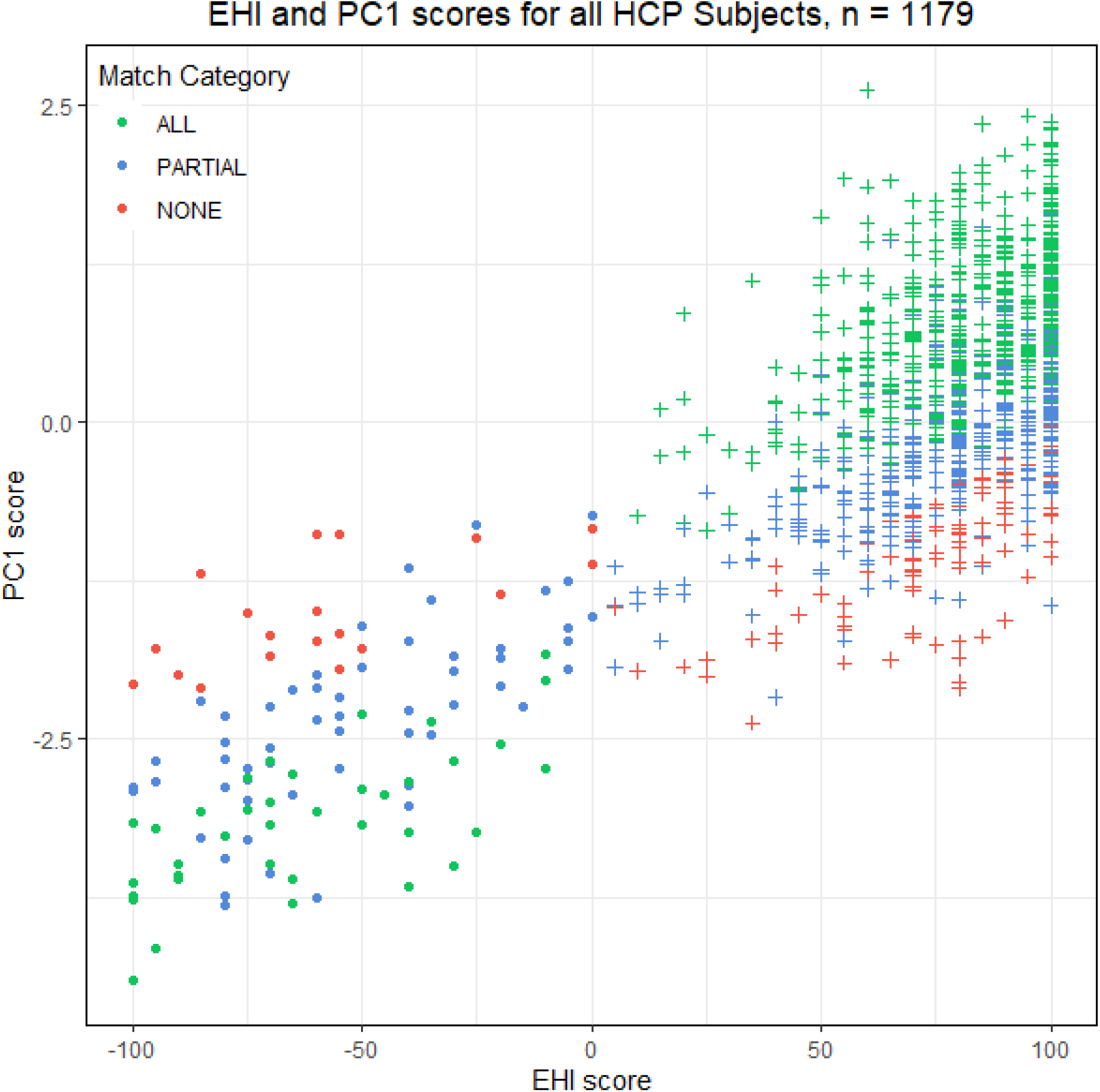
EHI and PC1 scores showing congruency categories for all HCP participants. EHI– participants are circles, EHI+ participants are crosses. PC1 scores generally track with EHI scores, but they also incorporate congruence data, as many EHI+ participants with partial or non-matched behavioral scores (i.e., left-hand-biased grip strength and/or pegboard LI’s) have negative PC1 values.

We will focus the rest of our discussion primarily on PC1 as an overall handedness measure. As PC1 encompasses all three handedness measures, we explored the effects of using PC1 scores, *not* the EHI or other raw handedness measures, as our variable for grouping participants. We repeated many of the analyses conducted on EHI+ and EHI– subjects, instead on PC1+ (*n =* 510) and PC1– (*n =* 669) subjects. T-tests on the raw behavioral scores show significant differences between PC1+ and PC1– participants, and had higher effect sizes overall than those from the EHI-based analyses (Table 7; compare to Table 2). As PC1 is a reformulation of the EHI and behavioral LI’s, each of these measures also differs significantly between PC1+ and PC1– participants (Figure 5). As discussed in our section on handedness consistency, a higher percentage of EHI+ participants were considered consistent handers compared to EHI– participants. When split by PC1 scores rather than EHI scores, handedness consistency information is still salient (**χ**-squared for EHI and consistency = 125.51, *p* < 0.001), with significantly more consistent handers in the PC1+ group (*n* = 487, 72.8%), and significantly less consistent handers in the PC1– group (*n* = 205, 40.2%). These results provide support for grouping subjects based on PC1, and further suggest that it might be a more encompassing, and thus more appropriate characterization of handedness in HCP participants.

**Table 7:**
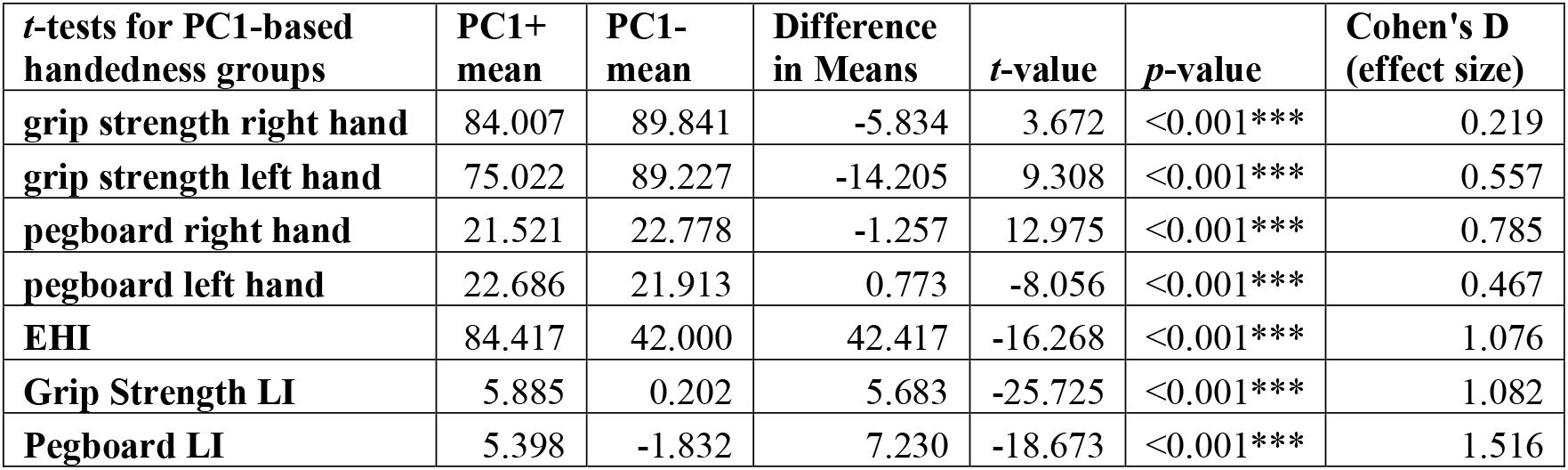
*t*-tests comparing PC1+ to PC1– means, for: raw grip strength for the right- and left-hands; raw pegboard times (in seconds) for the right- and left-hands; EHI values; and calculated grip strength and pegboard LI’s. Means, difference in means, t-values, p-values, and effect sizes are shown. * *p* < 0.05 ** *p* < 0.01 *** *p* < 0.001

| ***t*-tests for PC1-based handedness groups** | **PC1+ mean** | **PC1- mean** | **Difference in Means** | ***t*-value** | ***p*-value** | **Cohen's D (effect size)** |
| --- | --- | --- | --- | --- | --- | --- |
| **grip strength right hand** | 84.007 | 89.841 | -5.834 | 3.672 | <0.001*** | 0.219 |
| **grip strength left hand** | 75.022 | 89.227 | -14.205 | 9.308 | <0.001*** | 0.557 |
| **pegboard right hand** | 21.521 | 22.778 | -1.257 | 12.975 | <0.001*** | 0.785 |
| **pegboard left hand** | 22.686 | 21.913 | 0.773 | -8.056 | <0.001*** | 0.467 |
| **EHI** | 84.417 | 42.000 | 42.417 | -16.268 | <0.001*** | 1.076 |
| **Grip Strength LI** | 5.885 | 0.202 | 5.683 | -25.725 | <0.001*** | 1.082 |
| **Pegboard LI** | 5.398 | -1.832 | 7.230 | -18.673 | <0.001*** | 1.516 |

**Figure 5:**
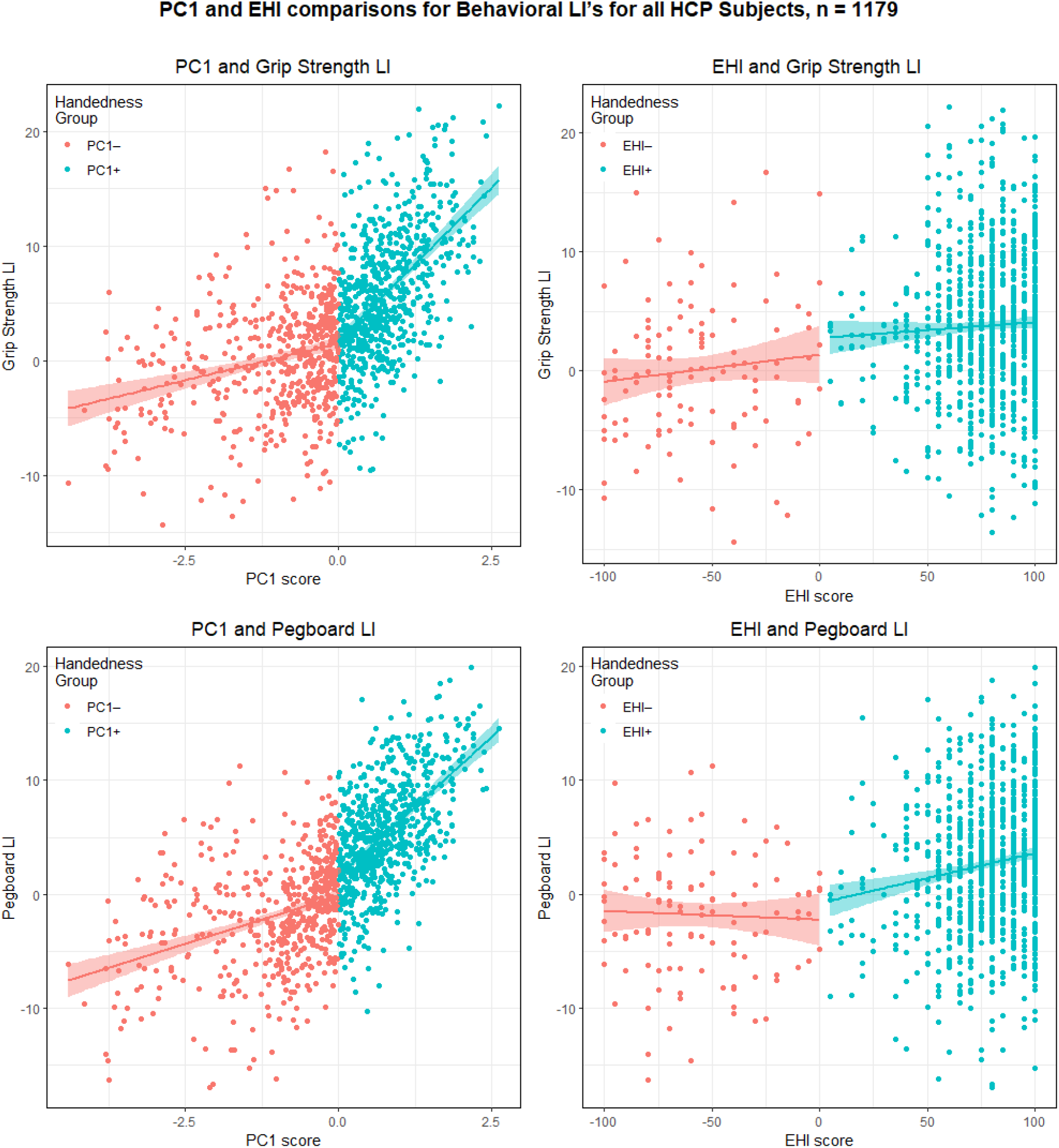
Comparison of behavioral handedness LI’s between PC1-based (left) and EHI-based (right) partitions of HCP subjects. EHI values showed poor correlations with grip strength and pegboard LI’s (see Table 3), whereas PC1-based groupings improve correspondence.

### Subject classification in neuroimaging contexts

In our final analysis, we explore how one might use PCA scores to address subject-specific disparities in their own data, in cases where multiple assessments of hand preference or performance were collected and a PCA can be run. As mentioned earlier, the HCP is a large-sample open neuroimaging initiative, so there is much that could be done in the future to explore how different handedness classification methods effect analyses on psychometric, anatomical, and functional brain data. Here, we use FreeSurfer brain segmentation data, available for direct download via the HCP, to illustrate the concept. Of the 1179 subjects with handedness data, 1092 (EHI+ *n* = 991, EHI– *n* = 101; PC1+ *n* = 510, PC1– *n* = 669) have cortical and subcortical segmentation data from the HCP FreeSurfer pipeline, which categorizes each voxel into one of over 30 cortical and subcortical classes (Fischl et al., 2002). FreeSurfer segmentations for each subject come from the HCP as whole-brain volumes (for example: white matter volume, sub-cortical grey matter volume, etc.), and as region-based surface areas, thicknesses, and volumes for the left- and right-hemispheres separately (for example, left-hemisphere *pars opercularis* area, right-hemisphere *pars triangularis* volume, etc.). In cases where paired left- and right-hemisphere data were given for a region, we calculated a laterality index (LI) for that region as before with the handedness measures, using the following formula:

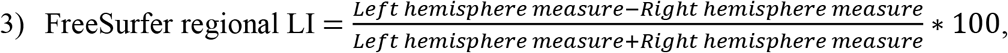

where positive values indicate a larger left-hemisphere measure, and negative values indicate a larger right-hemisphere measure.

This resulted in over 100 brain-based variables: 17 whole-brain volumes, volumetric LI’s for 16 regions, 34 surface area LI’s and 34 cortical thickness LI’s (see Supplementary Materials, Table 2 for a full list of Freesurfer regions).

Similar tests were run on the Freesurfer data (measured microliters (μl)) in as on the handedness metrics, although we applied Bonferroni corrections to these tests, as there were over 100 Freesurfer variables to test. T-tests were run on Freesurfer variables using subject groupings based on *both* the EHI and PC1 scores. For a majority of the Freesurfer measures, no significant differences were found between subject groups, regardless of whether they were defined by EHI scores or PC1 scores. However, many of the whole-brain volume measures were significantly different between PC1– and PC1+ subjects. After Bonferroni correction, Freesurfer data for the following regions showed significant differences between PC1+ and PC1– participants: Intracranial volume, Brain Segmentation volume, Cortical Grey Matter volume, Total Grey Matter volume, Total White Matter volume, and Supratentorial volume. Only Lateral Orbitofrontal area LI showed significant differences between EHI+ and EHI– participants, with all other comparisons resulting in nonsignificant corrected p-values (Table 8). Overall, we believe this provides tentative support for the use of PC1 scores as at least as good a dependent variable for subject handedness in future analyses as the EHI, although much more can be done to validate our approach with the other HCP neuroimaging data in the future. As many of these are raw measures, not LI’s, they do not directly relate to brain lateralization, but instead show that PC1– subjects in this sample have larger brains overall than PC1+ subjects (at α = 0.05). Linear models were run to explore the effects of age and sex on the Freesurfer findings, and in all cases, sex was the only significant predictor variable. Although there are similar proportions of males and females for the EHI-based grouping (**χ**-squared for EHI and sex = 2.612, *p* = 0.106), there are more females in the HCP sample overall (F = 638, 55%), and a significantly higher proportion of PC1+ females when the data are partitioned using PC1 scores (**χ**-squared for PC1 and sex = 32.707, *p* < 0.001). Interpretations for the Freesurfer data depend on further exploration of the relationship between handedness and sex in the HCP data, a fruitful avenue for future research.

**Table 8:** T-tests for Freesurfer data in PC1-based (left) and EHI-based (right) HCP data partitions. Only areas with significant Bonferroni-corrected *p*-values are shown (see Supplemental Data table 2 for a full list of Freesurfer regions). Bolded rows are significant at α = 0.05, and NS denotes non-significant *p*-values. * *p* < 0.05 ** *p* < 0.01 *** *p* < 0.001

| Whole Brain volumes | PC1 t-value | p-value | EHI t-value | p-value |
| --- | --- | --- | --- | --- |
| Intracranial volume | -3.923 | **0.009\*\*** | -0.095 | NS |
| Brain Segmentation volume | -4.162 | **0.003\*\*** | -0.363 | NS |
| Cortical Grey Matter volume | -4.102 | **0.004\*\*** | -0.285 | NS |
| Total Grey Matter volume | -4.391 | **0.001\*\*\*** | -0.331 | NS |
| Total White Matter volume | -3.466 | **0.050\*** | -0.210 | NS |
| Supratentorial volume | -3.974 | **0.008\*\*** | -0.334 | NS |
| Surface area LI's | t | p-value | t | p-value |
| Lateral Orbitofrontal area LI | -0.423 | NS | 2.049 | **0.043\*** |

## Discussion

In many ways, our results mirror previous investigations into the EHI as a measure of subject handedness in many aspects, but there are a few important differences. With a much larger sample size than previous studies (*n* = 1179), we show that several issues exist with the use of the EHI as a sole handedness metric. Based on our sub-sample analyses using the EHI to split subjects into groups, it seems that right- and left-handers have very different manifestations of multiple handedness measures. Future works on handedness-related asymmetries would benefit from whole-group *as well as* sub-sample reporting, as whole-sample analyses may be ‘benefitting’ from these disparities. For example, although the raw measures (grip strength and pegboard scores for the right and left hands) and LI’s had significant pair-wise correlations for the *whole sample*, our results corroborate other works in showing low correspondence in particular between EHI and grip strength LI, and we have also shown that—when considered separately—EHI, pegboard LI, and grip strength LI scores for EHI– subjects do not correlate with each other hardly at all. Although this not the case in some recent studies, many published works present analyses of their data for all subjects as one group, leaving us unsure of how much their effects are being driven by EHI+ subjects (especially in studies using non-balanced samples, such as our own).

The EHI no doubt communicates some credible information about subject handedness, but we have also shown that at least 10% of the entire HCP sample shows non-congruence between the survey- and behavioral-measures for handedness. Alarmingly, over 65 % of EHI– subjects have at least one directional disparity between EHI scores and Grip Strength and Pegboard LI’s. For those uninterested in studying handedness (those who purposely exclude ‘left-handers’ from their studies), these congruence results may also matter, as the most widely used criterion for left-hand subject exclusion in neuroimaging projects – a negative EHI score – is frequently not congruent with actual behavioral data, at least in the HCP subject pool. Regarding the pragmatics of screening for right-hand-exclusive paradigms, however, we believe the continued use of the EHI to be an appropriate course of action, particularly for participants with extreme EHI scores. However, for those explicitly studying handedness, our congruence data urge *extreme* caution for categorizing subjects based on EHI scores, particularly those with mid-range EHI values, and especially in studies specifically on handedness.

It is still generally unclear which handedness measures should be trusted more over the others in situations where they do not correlate, likely contributing to the continued use of only one measure, typically a survey-based one such as the EHI, in neuroimaging contexts. We have proposed an alternative approach for those who have previously collected multiple measures of handedness, which is to aggregate subject handedness data via Principal Components Analysis and use PC1 as an overall handedness metric (instead of EHI scores) as the independent variable for analyses. The benefit of this approach is that any number of handedness measures can be added to an analysis of this kind, which in turn may allow us to evaluate one measure against another in future studies. We have shown that many features of EHI-based analyses (correlations across measures, handedness consistency, etc.) are preserved when using PC1-based analyses, suggesting at the very least that this technique should be applied to additional data sets. A large caveat for this approach, however, is that data sets with different handedness measures may lead to very different PCA results and thus different PC1 values, than our specific example here. For example, if finger tapping were used instead of pegboard, it is not clear whether any of the resulting components would have similar loadings to our own, and so there might be a lack of an overall measure of handedness, as we have showed PC1 to be. The impact of different handedness measures on resulting PC’s should be further explored in future work on data sets both large and small.

In cases where EHI scores were the only handedness information collected, researchers can still contribute improve upon the discourse on handedness-related asymmetries by “reining in” their titles and text; instead of claiming that ‘*Handedness* does or does not correlate with X’ we might be more direct: ‘*EHI scores* do or do not correlate with X.’ After decades of conflicting results, it is time to seriously consider the value of the EHI as a sole proxy for subject handedness in lateralization studies. As Oldfield himself stated: “I am far from suggesting that, where manual or cerebral laterality are important issues in a piece of research, the [EHI] is a *sufficient* means of assessment of the handedness aspect. But for screening purposes…it may, I hope, prove useful” (1971: 110). We feel that researchers interested in laterality need to acknowledge that many of our problems in identifying handedness-related *and* handedness-independent asymmetries may be related largely to methods which are confounding, not clarifying, potentially real relationships. The best way to address this issue is to replicate works like those of Gonzalez and colleagues and use more naturalistic measures of handedness, even if they are labor-intensive. In lieu of this, and to make use of extant data such as that provided by the HCP and other open-source neuroimaging initiatives, it will also be important to compare each handedness measure to well-known asymmetries in the brain, perhaps starting with fMRI of language tasks, but extending beyond that. Careful evaluation of these measures may confirm the EHI as an important handedness measure to include in data analysis, but there is no sure way to know this without explicitly testing it.

## Acknowledgements

This research was supported in part by grant 52935 from the John Templeton Foundation titled: “What Drives Human Cognitive Evolution?”

## Data availability statement

The data which support the findings of this study are available from the Human Connectome Project (HCP). Some restrictions apply to the availability of these data, which were used under approval of the HCP for this study. Supporting data and R code for PCA calculations (for data like our own) can be made available from the corresponding author, LMR, under reasonable request. However, raw handedness measures, congruency data, and FreeSurfer values can *only* be provided when requests are accompanied by proof of prior permission for restricted data usage directly from the HCP (as detailed at https://www.humanconnectome.org/study/hcp-young-adult/document/restricted-data-usage).

## Footnotes

1 Some HCP data, including information about participant handedness, is considered “restricted” by the HCP, meaning that researchers must complete paperwork and request permission in order to gain access to that information. Information on restricted access for the HCP data can be found at https://www.humanconnectome.org/study/hcp-young-adult/document/restricted-data-usage

2 We had to ask for the raw NIH toolbox scores for the test-retest data, which lead us to an interesting discovery about the HCP’s definition of “raw” NIH toolbox data. It seems that the HCP “raw” NIH toolbox data for the 45 test-retest participants in the larger s1200 data release is actually *an average* of the two scores from T1 and T2, meaning that for these participants, our laterality indices (Grip strength LI and Pegboard LI) were computed on averaged “raw” data, unbeknownst to us, when the original analyses took place, whereas the remaining 1134 participant LI’s were calculated on single-measure (truly “raw”) data.

## References

Annett, M. (1985). Which theory fails? A reply to McManus. British Journal of Psychology, 76(1), 17–29. https://doi.org/10.1111/j.2044-8295.1985.tb01927.x

Badzakova-Trajkov, G, Corballis, MC, & Häberling, IS. (2016). Complementarity or independence of hemispheric specializations? A brief review. Neuropsychologia, 93, 386–393. https://doi.org/10.1016/J.NEUROPSYCHOLOGIA.2015.12.018

Bailey, L. M., McMillan, L. E., & Newman, A. J. (2019). A sinister subject: Quantifying handedness-based recruitment biases in current neuroimaging research. European Journal of Neuroscience, ejn.14542. https://doi.org/10.1111/ejn.14542

Barch, D. M., Burgess, G. C., Harms, M. P., Petersen, S. E., Schlaggar, B. L., Corbetta, M., … Van Essen, D. C. (2013). Function in the human connectome: Task-fMRI and individual differences in behavior. NeuroImage, 80, 1112–189. https://doi.org/10.1016/j.neuroimage.2013.05.033

Brown, S. G., Roy, E. A., Rohr, L. E., & Bryden, P. J. (2006). Using hand performance measures to predict handedness. Laterality: Asymmetries of Body, Brain and Cognition. https://doi.org/10.1080/1357650054200000440

Brown, S. G., Roy, E. A., Rohr, L. E., Snider, B. R., & Bryden, P. J. (2004). Preference and performance measures of handedness. Brain and Cognition, 55(2), 283–285. https://doi.org/10.1016/J.BANDC.2004.02.010

Bryden, M. P. (1977). Measuring handedness with questionnaires. Neuropsychologia, 15(4–5), 617–624. https://doi.org/10.1016/0028-3932(77)90067-7

Bryden, P. J., Brown, S. G., & Roy, E. A. (2011). Can an observational method of assessing hand preference be used to predict language lateralisation? Laterality: Asymmetries of Body, Brain and Cognition. https://doi.org/10.1080/1357650X.2010.513386

Büsch, D., Hagemann, N., & Bender, N. (2010). The dimensionality of the Edinburgh Handedness Inventory: An analysis with models of the item response theory. Laterality: Asymmetries of Body, Brain and Cognition, 15(6), 610–628. https://doi.org/10.1080/13576500903081806

Christman, S. D., Prichard, E. C., & Corser, R. (2015). Factor analysis of the Edinburgh Handedness Inventory: Inconsistent handedness yields a two-factor solution. Brain and Cognition, 98, 82–86. https://doi.org/10.1016/J.BANDC.2015.06.005

Corey, D. M., Hurley, M. M., & Foundas, A. L. (). Right and left handedness defined: a multivariate approach using hand preference and hand performance measures. Neuropsychiatry, Neuropsychology, and Behavioral Neurology, 14(3), 144–52. Retrieved from http://www.ncbi.nlm.nih.gov/pubmed/11513097

Dragovic, M. (2004). Towards an improved measure of the Edinburgh Handedness Inventory: A one-factor congeneric measurement model using confirmatory factor analysis. Laterality: Asymmetries of Body, Brain and Cognition, 9(4), 411–419. https://doi.org/10.1080/13576500342000248

Edlin, J. M., Leppanen, M. L., Fain, R. J., Hackländer, R. P., Hanaver-Torrez, S. D., & Lyle, K. B. (2015). On the use (and misuse?) of the Edinburgh Handedness Inventory. Brain and Cognition, 94, 44–51. https://doi.org/10.1016/J.BANDC.2015.01.003

Fagerland, M. W. (2012). t-tests, non-parametric tests, and large studies-a paradox of statistical practice? BMC Medical Research Methodology 12, 78. https://doi.org/10.1186/1471-2288-12-78

Fazio, R. L., & Cantor, J. M. (2015). Factor Structure of the Edinburgh Handedness Inventory Versus the Fazio Laterality Inventory in a Population With Established Atypical Handedness. Applied Neuropsychology: Adult, 22(2), 156–160. https://doi.org/10.1080/23279095.2014.940043

Fazio, R., Coenen, C., & Denney, R. L. (2012). The original instructions for the Edinburgh Handedness Inventory are misunderstood by a majority of participants. Laterality: Asymmetries of Body, Brain and Cognition, 17(1), 70–77. https://doi.org/10.1080/1357650X.2010.532801

Fazio, R., Dunham, K. J., Griswold, S., & Denney, R. L. (2013). An Improved Measure of Handedness: The Fazio Laterality Inventory. Applied Neuropsychology, 1–6. https://doi.org/10.1080/09084282.2012.684115

Fischl, B., Salat, D. H., Busa, E., Albert, M., Dieterich, M., Haselgrove, C., van der Kouwe, A., Killiany, R., Kennedy, D., Klaveness, S., Montillo, A., Makris, N., Rosen, B., Dale, A. M., (2002). Whole brain segmentation: automated labeling of neuroanatomical structures in the human brain. Neuron 33, 341–355. http://www.ncbi.nlm.nih.gov/pubmed/11832223

Flindall, J. W., & Gonzalez, C. L. R. (2018). Wait wait, don’t tell me: Handedness questionnaires do not predict hand preference for grasping. Laterality: Asymmetries of Body, Brain and Cognition, 1–21. https://doi.org/10.1080/1357650X.2018.1494184

Gershon, R. C., Cella, D., Fox, N. A., Havlik, R. J., Hendrie, H. C., & Wagster, M. V. (2010). Assessment of neurological and behavioural function: the NIH Toolbox. The Lancet Neurology, 9(2), 138–139. https://doi.org/10.1016/S1474-4422(09)70335-7

Gonzalez, C. L. R., & Goodale, M. A. (2009). Hand preference for precision grasping predicts language lateralization. Neuropsychologia, 47(14), 3182–9. https://doi.org/10.1016/j.neuropsychologia.2009.07.019

Gonzalez, C. L.R., van Rootselaar, N. A., & Gibb R. L. (2018). Sensorimotor lateralization scaffolds cognitive specialization. Progress in brain research 238: 405–433.

Gonzalez, C. L. R., Whitwell, R. L., Morrissey, B., Ganel, T., & Goodale, M. A. (2007). Left handedness does not extend to visually guided precision grasping. Experimental Brain Research, 182(2), 275–9. https://doi.org/10.1007/s00221-007-1090-1

Hopkins, B. (2018). A review of performance asymmetries in hand skill in nonhuman primates with a special emphasis on chimpanzees. Progress in brain research 238: 57–90.

Kallen, M., Slotkin, J., Griffinth, J., Magasi, M., Salsman, J., Nowinski, C., & Gershon, R. (2012a). NIH Toolbox 9-Hole Pegboard Dexterity Task. NIH Toolbox Technical Manual. Available from http://www.healthmeasures.net/images/nihtoolbox/Technical_Manuals/Motor/Toolbox_9-Hole_Pegboard_Dexterity_Test_Technical_Manual.pdf

Kallen, M., Slotkin, J., Griffinth, J., Magasi, M., Salsman, J., Nowinski, C., & Gershon, R. (2012b). NIH Toolbox Grip Strength Test. NIH Toolbox Technical Manual. Available from http://www.healthmeasures.net/images/nihtoolbox/Technical_Manuals/Motor/Toolbox_Grip_Strength_Test_Technical_Manual.pdf

Kaploun, K. A., & Abeare, C. A. (2010). A comparison of four handedness classification schemes through the investigation of lateralised semantic priming. Laterality: Asymmetries of Body, Brain and Cognition, 15(5), 481–500. https://doi.org/10.1080/13576500902958871

Koo, T. K., & Li, M. Y. (2016). A Guideline of Selecting and Reporting Intraclass Correlation Coefficients for Reliability Research. Journal of Chiropractic Medicine, 15(2), 155–63. https://doi.org/10.1016/j.jcm.2016.02.012

Króliczak, G., Gonzalez, C. L., & Carey, D. P. (2019). Editorial: Manual Skills, Handedness, and the Organization of Language in the Brain. Frontiers in Psychology. 10:930. https://doi.org/10.3389/fpsyg.2019.00930

Leppanen, M. L., Lyle, K. B., Edlin, F. M., & Schäfke, V. D. (2018). Is self-report a valid measure of unimanual object-based task performance? Laterality: Asymmetries of Body, Brain and Cognition, 23: 1–21. https://doi.org/10.1080/1357650X.2018.1550493

Mazoyer, B, Zago, L, Jobard, G, Crivello, F, Joliot, et al. (2014). Gaussian Mixture Modeling of Hemispheric Lateralization for Language in a Large Sample of Healthy Individuals Balanced for Handedness. PLoS ONE, 9(6), e101165. https://doi.org/10.1371/journal.pone.0101165

McManus, I. C. (1983). The Interpretation of Laterality. Cortex, 19(2), 187–214. https://doi.org/10.1016/S0010-9452(83)80014-8

McManus, I. C. (1984). The Power of a Procedure for Detecting Mixture Distributions in Laterality Data. Cortex, 20(3), 421–426. https://doi.org/10.1016/S0010-9452(84)80010-6

McManus, I. C. (1985). Right- and left-hand skill: Failure of the right shift model. British Journal of Psychology, 76(1), 1–16. https://doi.org/10.1111/j.2044-8295.1985.tb01926.x

McManus, I. C., Van Horn, J. D., & Bryden, P. J. (2016). The Tapley and Bryden test of performance differences between the hands: The original data, newer data, and the relation to pegboard and other tasks. Laterality: Asymmetries of Body, Brain and Cognition, 21(4–6), 371–396. https://doi.org/10.1080/1357650X.2016.1141916

Mellet, E, Jobard, G, Zago, L, Crivello, F, Petit, L, et al. (2014). Relationships between hand laterality and verbal and spatial skills in 436 healthy adults balanced for handedness. Laterality: Asymmetries of Body, Brain and Cognition, 19(4), 383–404.

Milenkovic, S., & Dragovic, M. (2013). Modification of the Edinburgh Handedness Inventory: A replication study, 18(3), 340–348. https://doi.org/10.1080/1357650X.2012.683196

Newman, S. D., Malaia, E., & Seo, R. (2014). Does degree of handedness in a group of right-handed individuals affect language comprehension? Brain and Cognition, 86, 98–103.

Oldfield, R. C. (1971). The assessment and analysis of handedness: the Edinburgh inventory. Neuropsychologia, 9(1), 97–113. Retrieved from http://www.ncbi.nlm.nih.gov/pubmed/5146491

Pritchard, E., Propper, R. E., & Christman, S. D. (2013). Degree of handedness, but not direction, is a systematic predictor of cognitive performance. Frontiers in Psychology 4.

R Core Team (2018). R: A Language and Environment for Statistical Computing. R Foundation for Statistical Computing, Vienna, Austria. https://www.R-project.org/

Reuben, D. B., Magasi, S., McCreath, H. E., Bohannon, R. W., Wang, Y.-C., Bubela, D. J., … Gershon, R. C. (2013). Motor assessment using the NIH Toolbox. Neurology, 80(11 Suppl 3), S65–75. https://doi.org/10.1212/WNL.0b013e3182872e01

Somers, M., Aukes, M. F., Ophoff, R. A., Boks, M. P., Fleer, W., de Visser, K. (C). L., … Sommer, I. E. (2015). On the relationship between degree of hand-preference and degree of language lateralization. Brain and Language. https://doi.org/10.1016/j.bandl.2015.03.006

Tapley, S. M., & Bryden, M. P. (1985). A group test for the assessment of performance between the hands. Neuropsychologia, 23(2), 215–221. https://doi.org/10.1016/0028-3932(85)90105-8

Veale, J. F. (2014). Edinburgh Handedness Inventory – Short Form: A revised version based on confirmatory factor analysis. Laterality: Asymmetries of Body, Brain and Cognition, 19(2), 164–177. https://doi.org/10.1080/1357650X.2013.783045

Van Essen, D. C., Smith, S. M., Barch, D. M., Behrens, T. E. J., Yacoub, E., Ugurbil, K., & WU-Minn HCP Consortium (2013). The WU-Minn Human Connectome Project: an overview. NeuroImage, 80, 62–79. https://doi.org/10.1016/j.neuroimage.2013.05.041

Van Horn, J. D. (1992). Brain structural abnormality and laterality in schizophrenia (Unpublished PhD thesis). University College London.

Vingerhoets, G. (2014). Praxis, language, and handedness: a tricky triad. Cortex 57, 294–296. doi: 10.1016/j.cortex.2014.01.019

Wang, Y. C., Magasi, S. R., Bohannon, R. W., Reuben, D. B., McCreath, H. E., Bubela, D. J., Gershon, R. C., & Rymer, W. Z. (2011). Assessing dexterity function: A comparison of two alternatives for the NIH toolbox. Journal of Hand Therapy, 24(4), 313–321. https://doi.org/10.1016/j.jht.2011.05.001

Willems, R. M., Der Haegen, L. Van, Fisher, S. E., & Francks, C. (2014). On the other hand: Including left-handers in cognitive neuroscience and neurogenetics. Nature Reviews Neuroscience. https://doi.org/10.1038/nrn3679

Zago, L, Petit, L, Mellet, E, Jobard, G, Crivello, F, et al. (2016). The association between hemispheric specialization for language production and for spatial attention depends on left-hand preference strength. Neuropsychologia, 93, 394–406. https://doi.org/10.1016/j.neuropsychologia.2015.11.018

